# Sequence variability of BamA and FadL candidate vaccinogens suggests divergent evolutionary paths of *Treponema pallidum* outer membrane proteins

**DOI:** 10.1101/2025.04.21.649848

**Authors:** Everton B. Bettin, Farhang Aghakhanian, Christopher M. Hennelly, Wentao Chen, Timothy C. Davenport, Simon T. Hackl, Andre A. Grassmann, Fabio Vargas-Cely, Sebastián Silva, Jonny A. García-Luna, Lady G. Ramirez, Yinbo Jiang, Ligang Yang, Heping Zheng, Bin Yang, Petra Pospíšilová, David Šmajs, Mitch M. Matoga, Irving F. Hoffman, Eduardo López-Medina, Kay Nieselt, M. Anthony Moody, Arlene C. Seña, Juan C. Salazar, Jonathan B. Parr, Melissa J. Caimano, Kelly L. Hawley, Justin D. Radolf

## Abstract

Knowledge of *Treponema pallidum* subspecies *pallidum* (*TPA*) outer membrane protein (OMP) sequence variability is essential for understanding spirochete proliferation within endemic populations as well as design of a globally effective syphilis vaccine. Our group has identified extracellular loops (ECLs) of *TPA* BamA (TP0326) and members of the FadL family (TP0548, TP0856, TP0858, TP0859, and TP0865) as potential components of a multivalent vaccine cocktail. As part of a consortium to explore *TPA* strain diversity, we mapped the variability of BamA and FadL orthologs in 186 *TPA* strains from Malawi, China and Colombia onto predicted 3D structures. The 186 genomes fell into eight subclades (five Nichols-, three SS14-lineage) with substantial geographic restriction. Single nucleotide variants accounted for the large majority of proteoforms, with variability notably higher within the Nichols-lineage strains. Most mutations were in regions of the proteins predicted to be extracellular and harboring B cell epitopes. We observed a striking difference in the degree of variability between the six OMPs, suggesting that these proteins are following divergent evolutionary paths. Concatenation of OMP sequences recapitulated the phylogenetic structure of the *TPA* strains, effectively segregating within clades and largely clustering by subclades. Lastly, we noted that BamA and FadL candidate ECL vaccinogens previously shown to elicit antibodies that kill treponemes during *in vitro* cultivation are well conserved. Taken as a whole, our study establishes a structural-phylogenetic approach for analyzing the forces shaping the host-pathogen interface in syphilis within endemic populations while informing selection of vaccine targets.

**IMPORTANCE:** Syphilis remains a major global health concern, reinforcing the need for a safe and effective vaccine. Understanding the variability of *TPA* OMPs is essential for tracking pathogen evolution and informing vaccine design. Here, we analyzed the variability of six *TPA* OMPs in 186 strains from Malawi, China, and Colombia, identifying protein-specific evolutionary patterns. Most mutations were localized in extracellular regions and, notably, appeared to correlate with the phylogenetic structure of *TPA*. Despite OMP heterogeneity, several candidate vaccinogens remained highly conserved, reinforcing their potential as globally effective vaccine targets. Our study establishes a structural-phylogenetic framework for dissecting the forces shaping the host-spirochete interface within endemic populations and provides a foundation for designing a globally effective syphilis vaccine.

## INTRODUCTION

Syphilis is a multistage, sexually transmitted infection caused by the highly invasive and immunoevasive spirochete *Treponema pallidum* subspecies *pallidum* (*TPA*)(1, 2). The disease has undergone a worldwide resurgence since the start of the new millennium(3); even though its causative agent remains exquisitely susceptible to penicillin after more than seven decades of use(3, 4). These alarming epidemiological trends underscore the need for a safe and effective vaccine as well as an improved understanding of the basic biology and virulence properties that foster proliferation of the syphilis spirochete within at-risk populations(5–7). *TPA* is a diderm bacterium with an unorthodox outer membrane (OM) lacking lipopolysaccharide (LPS) and containing a low density of integral outer membrane proteins (OMPs) and a paucity of surface-exposed lipoproteins(5, 8–10). The molecular architecture of the spirochete’s OM is the ultrastructural basis for its impressive capacity to evade innate and adaptive host defenses, giving rise to its designation as the ‘stealth pathogen’(5, 10, 11). Despite the lack of sequence similarity between *TPA* OMPs and their counterparts in Gram-negative bacteria, advances in protein three-dimensional (3D) structure prediction have enabled the identification of transmembrane β-barrel-forming proteins encoded within the *TPA* genome(10, 12). *TPA*’s repertoire of OMPs (the *TPA* ‘OMPeome’) includes two stand-alone proteins, BamA and LptD, involved in OM biogenesis and four paralogous families involved in importation of nutrients or extrusion of noxious substances across the OM: 8-stranded β-barrels, OM factors for efflux pumps, *TPA* repeat proteins, and orthologs for FadL long-chain fatty acid transporters(10, 13).

*TPA* is widely considered to be an extracellular bacterium(5, 14, 15). To effect spirochete clearance, antibodies elicited during infection or by vaccination must target extracellular regions of the bacterium’s OMPs(5–7). In recent years, we have identified surface-exposed regions in BamA and in members of the FadL family as candidate components of an experimental syphilis vaccine cocktail(16–19). BamA (TP0326) is the central component of the molecular machinery that inserts newly exported OMPs into the spirochete’s OM(20). Like its Gram-negative counterparts, BamA in *TPA* is bipartite and consists of a predicted 16-stranded β-barrel with eight extracellular loops (ECLs) and five polypeptide transport-associated (POTRA) domains within the periplasm(13, 17, 21). The five *TPA* FadL-like orthologs (TP0548, TP0856, TP0858, TP0859, and TP0865) contain structural features found in canonical fatty acid transporters: a 14-stranded β-barrel with seven ECLs and an N-terminal hatch domain that plugs the lumen of the barrel(22, 23). Unlike canonical FadLs, however, the hatches of the *TPA* orthologs are predicted to extend into the extracellular milieu where they should be antibody accessible(13). Three *TPA* FadLs (TP0548, TP0859, and TP0865) also contain unique C-terminal tetratricopeptide (TPR) domains predicted to reside in the periplasmic compartment(13, 24).

Knowledge of *TPA* OMP variability is a prerequisite for understanding spirochete proliferation within endemic regions as well as the design of a globally efficacious syphilis vaccine. While advancements in whole-genome sequencing (WGS) of *TPA* strains in clinical specimens have significantly expanded our knowledge of *TPA* genetic diversity(25–34), these analyses generally have not focused upon OMP sequence variability. Moreover, publicly available *TPA* genomic sequences usually represent a limited number of strains from different geographic locations as opposed to deep sampling at individual sites. Herein, as part of a global consortium to explore *TPA* strain diversity in the context of syphilis vaccine design(31), we sequenced 186 *TPA* strains from our clinical research sites in Malawi, China, and Colombia, and subsequently mapped the variability of BamA and the five FadL orthologs onto their predicted 3D structures. Most mutations were in regions predicted to be extracellular and harboring B cell epitopes, suggesting that host immune pressure is a major driver of OMP diversity. A striking difference in the degree of variability between the six OMPs was observed, suggesting they are following divergent evolutionary paths. The OMP profiles generated for each strain within our cohort recapitulated the phylogenetic structure of *TPA*, segregating by clades and largely by subclades. These findings indicate that recombination is not a major factor underlying variation of this portion of the OMPeome, and they raise the possibility that pressures arising from demographic or regional factors contribute to OMP evolution. Lastly, we noted a high degree of conservation among BamA and FadL candidate ECL vaccinogens that elicit antibodies capable of killing treponemes during *in vitro* cultivation(18, 19). Taken as a whole, our study establishes a structural-phylogenetic approach for analyzing the forces shaping the host-spirochete interface within endemic populations. Our study also informs the selection of surface-exposed targets for a vaccine that could eliminate a disease that has afflicted humankind for centuries.

## RESULTS

### Sources and phylogenies of *TPA* genomes

To explore *TPA* strain diversity in the context of syphilis vaccine design(31), we recruited individuals with early syphilis (primary, secondary and early latent syphilis) at clinical research sites in Lilongwe, Malawi; Guangzhou, China; and in Cali, Colombia. Genomic sequences from *TPA* strains infecting 131 patients were recently reported(31). For the present study, we added 55 unique genomes from all three sites (Figure 1A), yielding a total of 186 genomic sequences obtained between 2015 and 2022 (see Supplementary Table 1 for strain information).

**Figure 1.**
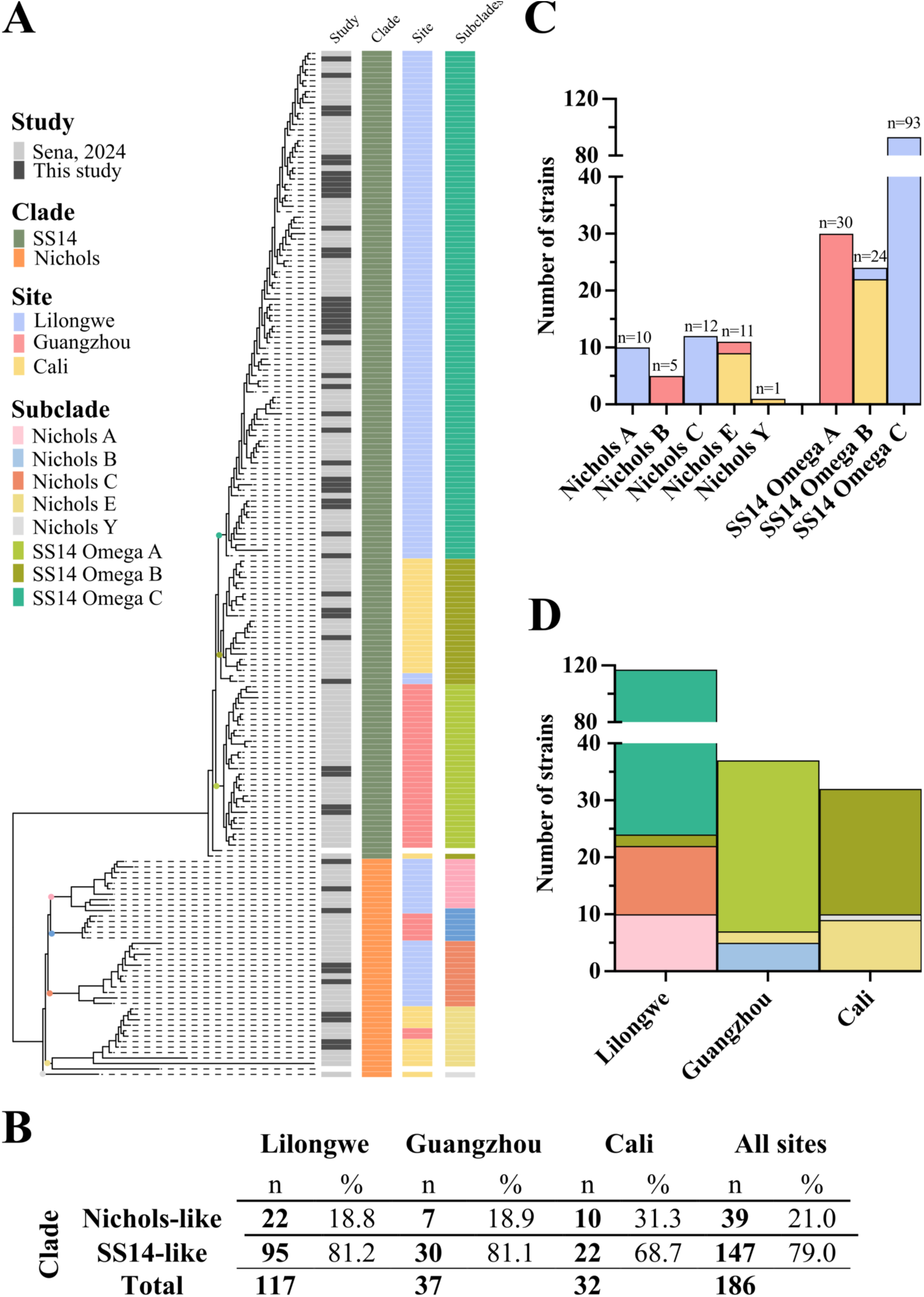
Distribution of the 186 strains within our cohort according to their subclades and sites. **(A)** Recombination-masked *TPA* whole-genome phylogeny demonstrated that SS14-lineage strains were predominant within our cohort. Strains were assigned to eight distinct *TPA* subclades: five Nichols (Nichols A, B, C, E, and Y) and three SS14 (SS14 Omega A, B, and C)**. (B)** Similar proportions of Nichols- and SS14-lineage strains were found in Lilongwe and Guangzhou (∼81.1%), but a lower (68.7%) in Cali. **(C and D)** All *TPA* subclades exhibited a high degree of geographic structure.

While the majority (∼78.5%) of strains were directly sequenced from genital ulcer swabs or secondary syphilis skin biopsies, 37 genomic sequences from Guangzhou and 3 from Cali were obtained after *TPA* passage in rabbits (Supplementary Table 1). We have previously confirmed that rabbit passage has minimal effects on *TPA* genomic stability and does not introduce mutations in the analyzed loci(32). In accord with other recent studies(25–28, 33), phylogenomic analysis revealed a predominance (79%; n=147) of SS14-lineage strains within our cohort (Figure 1A and 1B). The proportion of SS14-lineage strains was virtually identical (∼81%) in Lilongwe and Guangzhou but, lower (68%) in Cali (Figure 1B); of note, a higher prevalence of Nichols-lineage strains has been observed in other genomic surveys from South America(26, 28).

To facilitate comparison with previous genomic analyses(31, 35), the strains were curated based on maximum likelihood phylogeny (Figure 1A), resulting in five Nichols (Nichols A, B, C, E and Y) and three SS14 (SS14 Omega A, B and C) subclades. The subclade formerly called SS14 Omega East-Asia(31, 35) herein was renamed SS14 Omega A. The remaining SS14-lineage strains, previously grouped as a single SS14 Omega subclade(31), formed two distinct branches on the tree, and, therefore, were classified as part of the SS14 Omega B and C subclades, respectively (Figure 1A). All subclades exhibited a substantial degree of geographic restriction (Figure 1C), with only one lineage in each clade (Nichols E and SS14 Omega B) containing a small number of strains from another site. Also noteworthy is that different combinations of Nichols and SS14 subclades were co-circulating in each site. All three sites contained strains from two Nichols subclades (Figure 1D), although Lilongwe, which provided the most strains to the cohort, was the only site containing strains from two different SS14 subclades (Omega B and C).

### *TPA* BamA and FadL 3D models

Using a Biopython-based computational framework developed by our group, we identified mutations in the BamA and FadL sequences within our cohort and mapped them onto 3D structures generated by AlphaFold3(36). All β-strand residues exhibited high confidence values at a minimum, as defined by Alphafold3, supporting accurate delineation of the boundaries between β-strands and ECLs (Supplementary Figure 1). Notably, in the AlphaFold3 model for BamA, ECL3 is longer and ECL4 is shorter than previously predicted by the ModWeb server, which used the *Neisseria gonorrhoeae* BamA (PDB ID: 4K3B) as a template(17) (Supplementary Figure 2). AlphaFold3 models for the FadLs aligned closely with those previously generated using trRosetta(13). Both algorithms predict that the *TPA* FadLs lack a β-strand three kink, a characteristic feature of canonical FadLs(23, 37), and have hatches that extend into the extracellular milieu, suggesting that *TPA* orthologs may use alternative mechanisms for substrate capture and transport.

### Substitutions in BamA occurred exclusively in ECLs and largely segregated with subclades

BamA in the Nichols and SS14 reference strains (designated proteoforms 1 and 2, respectively) differ by nine residues with six of these mutations in ECL3 (Figure 2). Substitutions in five of these nine reference strain ‘discriminators’ are non-conservative, as determined by Grantham physicochemical distance(38). Three of the reference strain discriminators (T543, R545, and F656 in the Nichols reference) were conserved throughout the cohort. We identified nine Nichols variants containing 15 intra-clade mutations (proteoforms 3-11) and four SS14 variants with six intra-clade mutations (proteoforms 12-15); most replacements arose from single nucleotide variants (SNVs) (Supplementary Table 2). Only one of 39 Nichols-lineage strains (Nichols E from Cali) contained a BamA identical to the Nichols reference. In contrast, 53 of 147 strains, found predominantly in the SS14 Omega A and B subclades, contained exact matches to the SS14 reference sequence. All mutations were located exclusively in ECLs and largely segregated with subclades (*e.g*., variable ECLs 3 and 6 in Nichols A, ECL7 in Nichols E and ECL8 in SS14 Omega B and C). The vast majority (92 of 94) of the SS14-lineage strains in our cohort with a variant BamA were in the Omega C subclade and contained a G841D mutation in ECL8 (proteoform 14).

**Figure 2.**
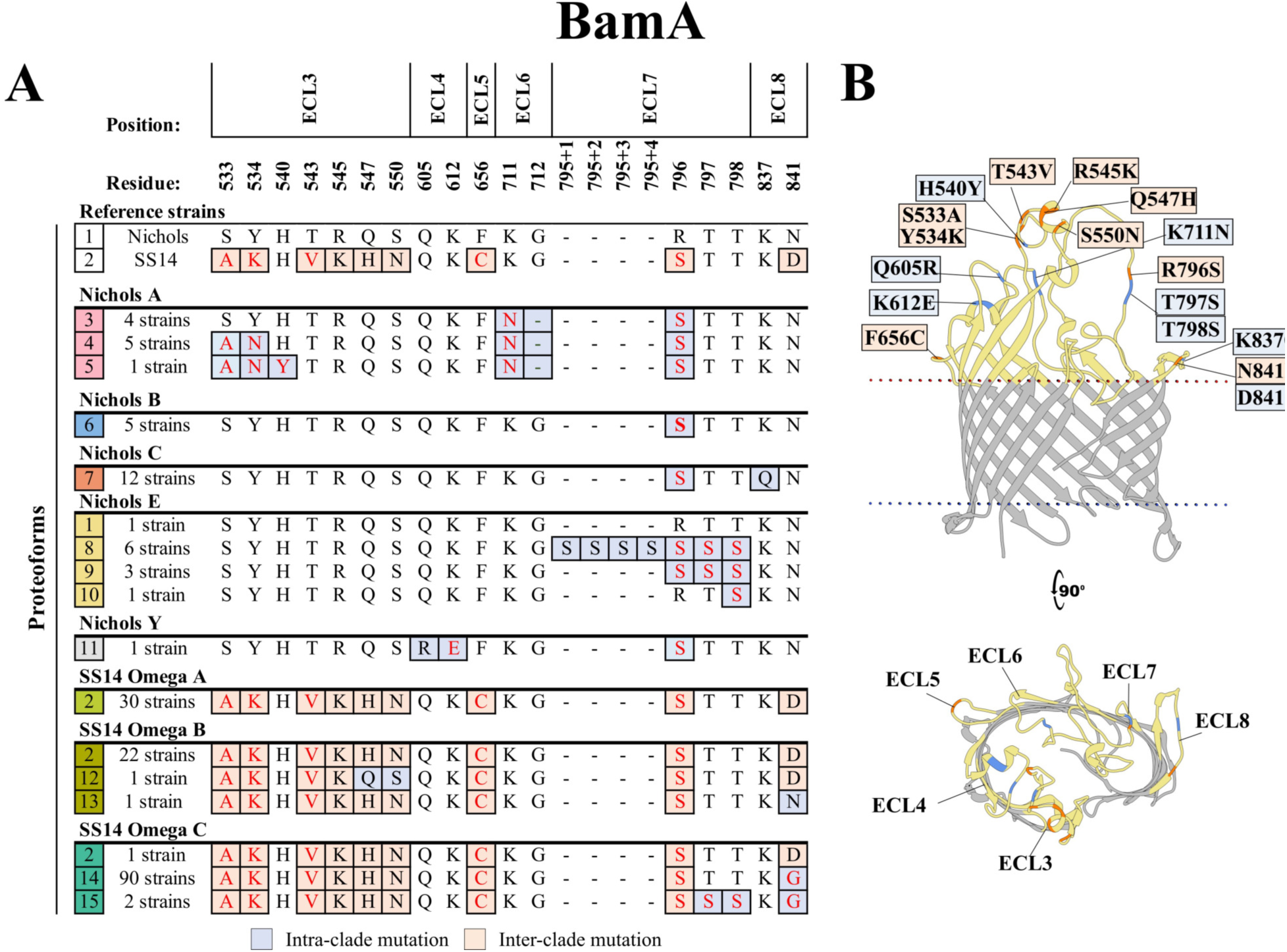
Substitutions in BamA are exclusively in ECLs and largely segregate with *TPA* subclades. **(A)** Amino acid residues differences between the 15 identified BamA proteoforms (1-15 in the leftmost squares). Nine 8discriminator9 residues (orange boxes) distinguish the proteoforms in Nichols and SS14 reference strains. We identified 17 additional intra-clade mutations (blue boxes) in ECLs. Non-conservative substitutions are colored in red. Residue are numbered according to their position in the Nichols reference sequence. **(B)** Amino acid substitutions were mapped onto the AlphaFold3 model for the Nichols reference BamA ³-barrel. Periplasmic POTRA domains are hidden in the model to facilitate visualization.

### TP0856 and TP0859 are highly conserved

The TP0856 sequences in Nichols and SS14 reference strains (proteoform 1) are identical. No mutations in TP0856 were observed in the 39 Nichols-lineage strains in our cohort (Figure 3A and 3B). Two SS14 Omega A strains from Guangzhou and one Omega B strain from Cali contained SNVs leading to non-conservative substitutions in ECL7 and hatch (proteoforms 2 and 3), respectively (Figure 3A and 3B, and Supplementary Table 2). Notably, two SS14 Omega C strains from Lilongwe harbored *tp0856* alleles containing a nonsense SNV at position 327 (Figure 3A and 3B, and Supplementary Table 2). TP0856 was the only OMP analyzed herein with more variants found within the SS14-lineage. As with TP0856, the Nichols and SS14 reference sequences for TP0859 are identical (proteoform 1) (Figure 3C and 3D). SNVs generated four TP0859 variant proteoforms in a small number of strains (seven Nichols A and five Omega C) from Lilongwe.

**Figure 3.**
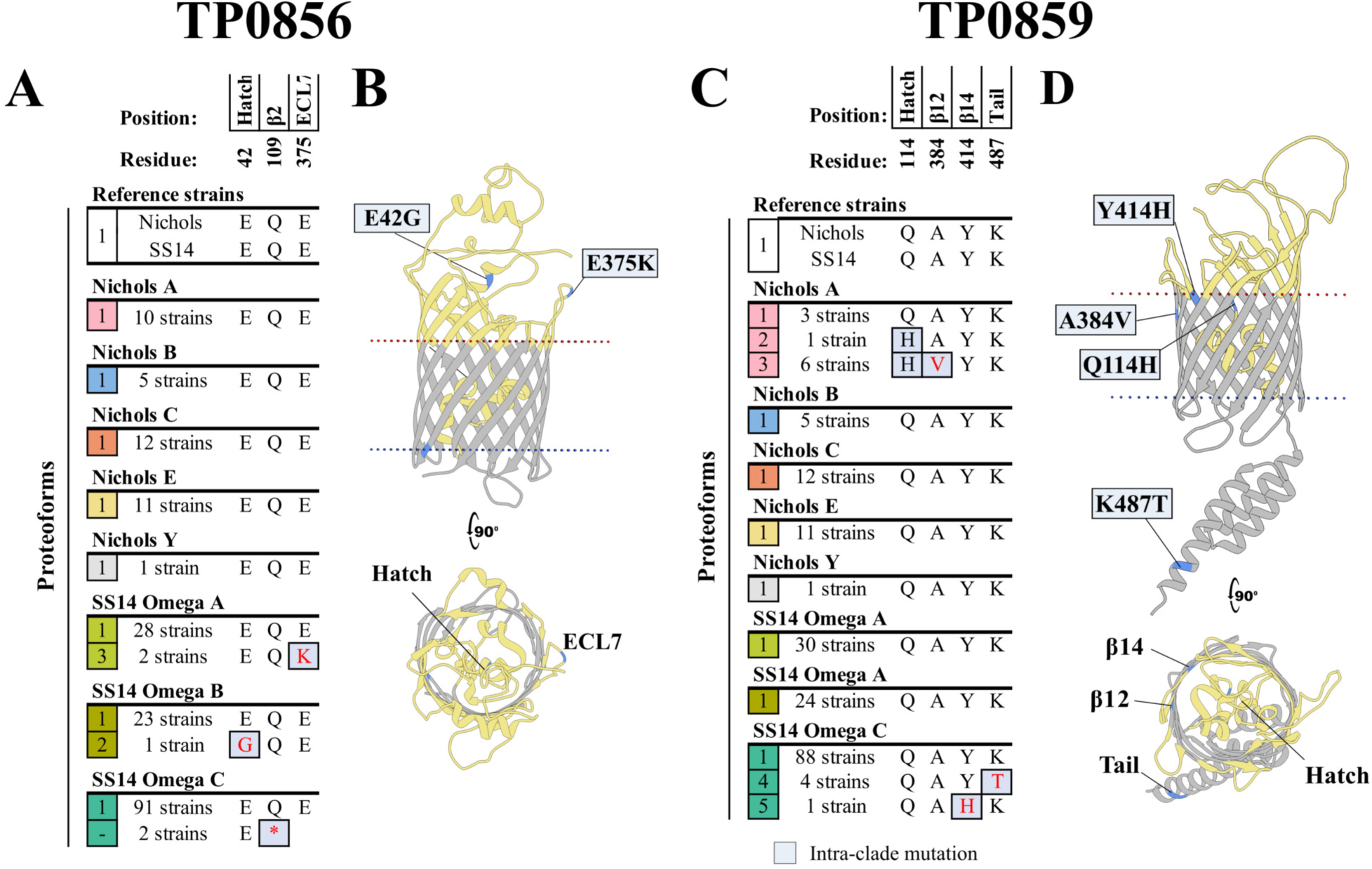
TP0856 and TP0859 are highly conserved. TP0856 and TP0859 proteoforms from Nichols and SS14 reference strains are identical. **(A)** Amino acid residues differences between the three identified TP0856 proteoforms (1-3 in the leftmost squares). Only three SS14-lineage strains contained TP0856 proteoforms with intra-clade mutations (blue boxes). Two *tp0856* sequences contained an early stop-codon at position 327. **(B)** Predicted mutations in ECL7 and the hatch of TP0856 variants from Guangzhou (SS14 Omega A) and Cali (Omega B). **(C)** Amino acid residues differences between the five identified TP0859 proteoforms (1-5 in the leftmost squares). Intra-clade mutations were identified in seven Nichols-lineage and five SS14-lineage strains. **(D)** Predicted mutations in the hatch, ³-strands, and periplasmic regions of TP0859 variants. A portion of the hatch is hidden in the model to facilitate visualization. In **A** and **C** non-conservative substitutions are colored in red. Residues are numbered according to their position in the Nichols reference sequence.

### TP0858 variants result primarily from intra-clade mutations but also a rare recombination

A single, conservative amino acid substitution due to a SNV in ECL7 (S380N) distinguishes the TP0858 reference sequences in Nichols and SS14 (proteoforms 1 and 2; Figure 4 and Supplementary Table 2). Intra-clade substitutions generated by SNVs at seven different positions, along with a recombination in one strain, gave rise to four TP0858 variants in each clade (proteoforms 3-6 and 7-10, respectively). More than half (24 of 39) of the Nichols-lineage strains contained TP0858 variants compared to only 12 of the 147 SS14-lineage strains. The sole Nichols Y strain contained a TP0858 variant (proteoform 6) with a previously reported (39, 40) recombination in ECL4 that replaces residues 276-282 with the corresponding residues (271–277) of TP0856. Interestingly, one SS14 Omega A strain harbored the Nichols reference due to a SNV that converted the single SS14 reference discriminator asparagine at position 380 to a serine residue.

**Figure 4.**
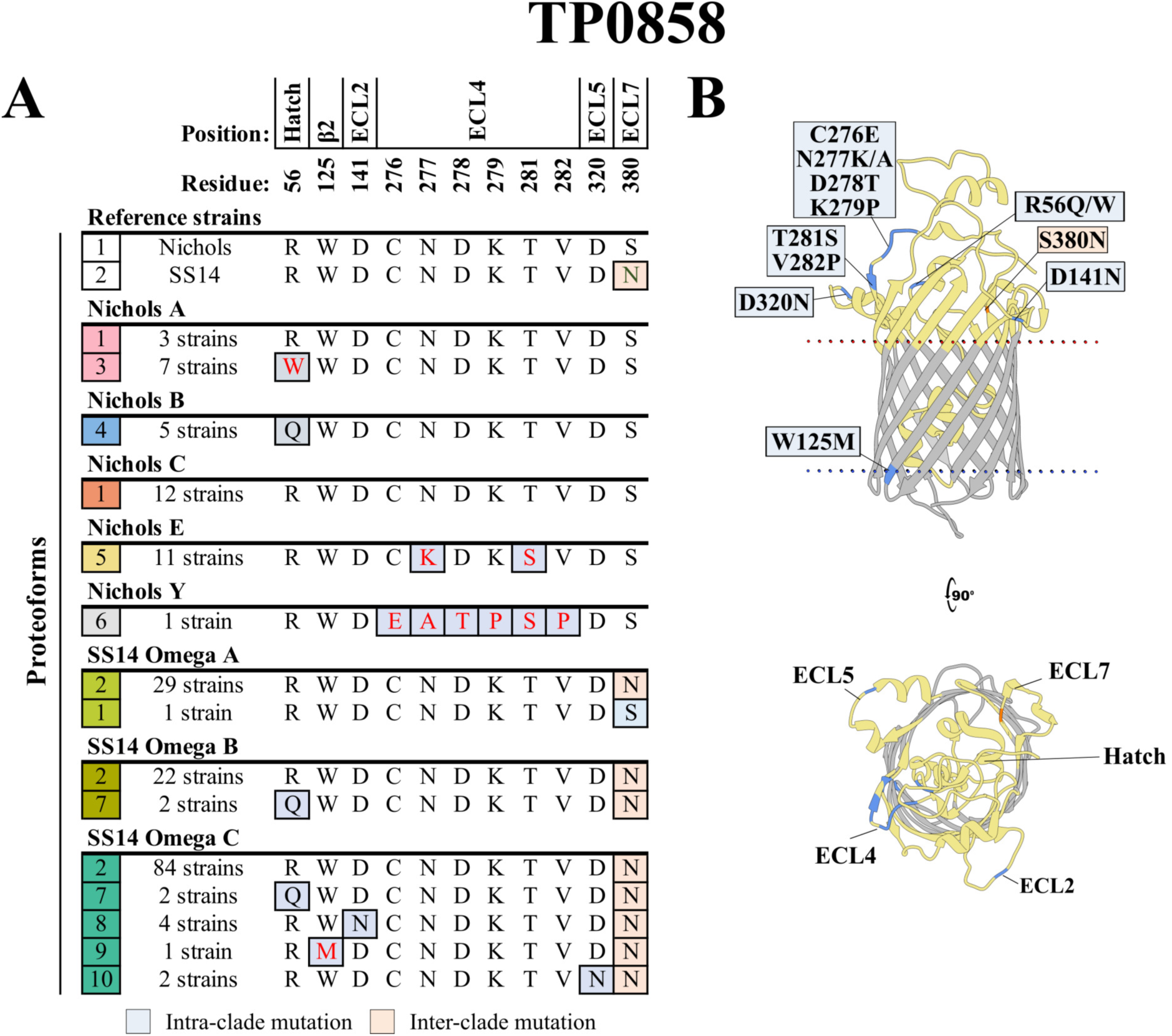
Amino acid substitutions in TP0858 result primarily from intra-clade mutations but also a rare recombination. **(A)** Amino acid residues differences between the 10 identified TP0858 proteoforms (1-10 in the leftmost squares). The TP0858 sequences in the Nichols and SS14 reference genomes differ by only a S380N substitution (orange boxes). Eight variant proteoforms containing intra-clade mutation (blue boxes) were identified within the cohort. All eleven Nichols E strains carried mutations (N277K and T281S) in ECL4. In the sole Nichols Y strain, a putative recombination between ECL4 of TP0856 and TP0858 was identified within residues 276-282 of ECL4. Non-conservative substitutions are colored in red. Residues are numbered according to their position in the Nichols reference sequence. **(B)** Substitutions mapped onto the AlphaFold3 model for the Nichols reference.

### TP0548 is hypervariable due to Nichols and SS14 clade-specific substitutions within the hatch and ECL2, respectively

The Nichols and SS14 TP0548 reference sequences (proteoforms 1 and 2) differ by 22 substitutions (17 non-conservative) along with a three amino acid insertion in ECL2 of the SS14 reference OMP (Figure 5). We identified a total of 16 variants containing intra-clade mutations generated by SNVs in 15 positions, eight at reference strain discriminators (Figure 5 and Supplementary Table 2). Within the Nichols clade, only one Nichols A strain (from Lilongwe) contained a sequence identical to the clade reference. A striking feature of TP0548 variants within the Nichols-lineage (proteoforms 3-9) was the predominance of intra-clade substitutions predicted to reside in the extracellular portion of the hatch. Among the 147 SS14-lineage strains, there were no exact matches to the reference due to a ubiquitous glycine insertion after position 51 of the hatch. A variant defined exclusively by this insertion (proteoform 10) was found in all 30 SS14 Omega A strains and in 18 of 24 Omega B strains, but in only one Omega C strain. Intriguingly, residues of the hatch that were variable within the Nichols clade were highly conserved across SS14-lineage strains. However, 92 of the SS14 Omega C strains contained substitutions within five closely spaced residues in ECL2 (proteoforms 12-18).

**Figure 5.**
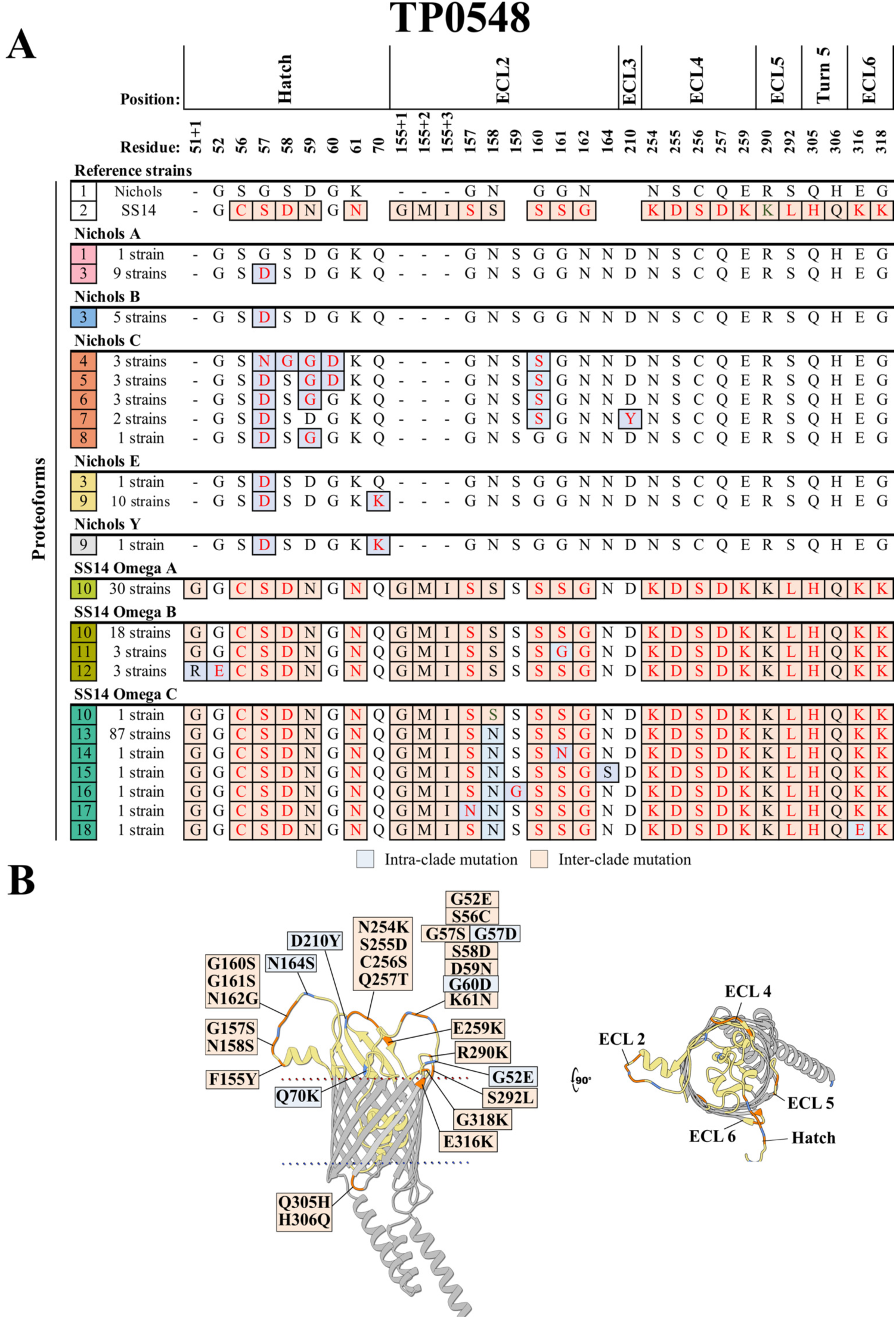
TP0548 is hypervariable with Nichols- and SS14-specific substitutions within the hatch and ECL2, respectively. **(A)** Amino acid residue differences between the 18 identified TP0548 proteoforms (1-18 in the leftmost squares). Reference sequences for Nichols and SS14 contain 25 inter-clade mutations (orange boxes). We identified 16 variant proteoforms within our cohort, eight in each clade, containing intra-clade substitutions (blue boxes) at 15 amino acid positions. Most of the intra-clade differences are found in the hatch and in ECL2, for the Nichols-lineage and SS14-lineage strains, respectively. Non-conservative substitutions are colored in red. Residues are numbered according to their position in the Nichols reference sequence. **(B)** Substitutions were mapped onto the AlphaFold3 model for the Nichols reference.

### TP0865 in the Nichols E subclade is hypervariable due to an inter-subspecies recombination

The Nichols and SS14 TP0865 reference sequences (proteoforms 1 and 2) differ by non-conservative and conservative amino acid substitutions originating, respectively, from SNVs within ECL2 (A193T) and β-strand 12 (S372N) along with the insertion of an asparagine after position 237 in ECL3 of the SS14 reference (Figure 6 and Supplementary Table 2). Only one reference strain discriminator (S372N) was conserved within all strains within the Nichols lineage, while the SS14-lineage strains contained no substitutions in discriminators. Only two of the 39 Nichols-lineage strains (Nichols C, Lilongwe and Nichols Y, Cali) contained TP0865 sequences identical to the clade reference. We identified five variants (proteoforms 3-7) within the Nichols A-C subclades containing substitutions at four positions within the hatch, ECL3 and ECL7. By contrast, the Nichols E subclade contained nine variants (proteoforms 8-16) harboring substitutions at 27 positions predominantly in the hatch and ECLs 2-4. The majority (19 of 27) of the substitutions were in two stretches of DNA encoding residues 89-193 and 289-395 (*i.e*., flanking ECL3) previously reported to have arisen by recombination between *TPA* and endemicum treponemes (*TEN/TPE*)(41–43) (Supplementary Figure 3); an additional eight substitutions generated hypervariability in the nonrecombinant stretch encoding ECL3. In stark contrast, only one of the 147 SS14-lineage strains (Omega C, Lilongwe) contained a non-conservative substitution (P454S) in the periplasmic TPR domain (proteoform 17).

**Figure 6.**
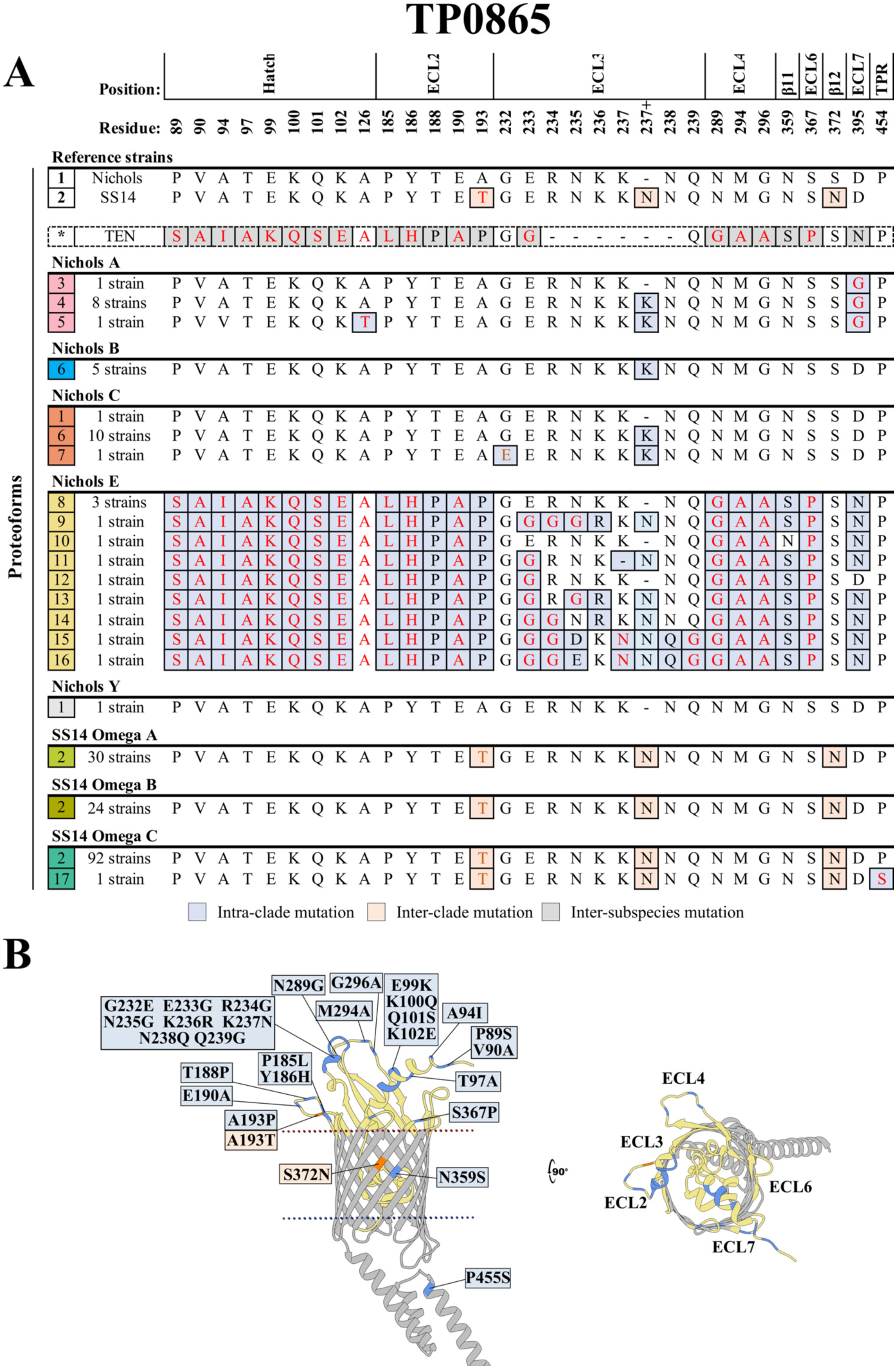
TP0865 in the Nichols E subclade contains an ancient inter-subspecies recombination and a hypervariable ECL3. **(A)** Amino acid residues differences between the 17 identified TP0865 proteoforms (1-17 in the leftmost squares). Three inter-clade mutations (orange boxes) distinguish the Nichols and SS14 references. We identified 15 variant TP0865 proteoforms containing intra-clade mutations (blue boxes), nine variants were within the Nichols E subclade. The high variability observed within Nichols E aligns with previous report of recombination events between *TPA* and endemicum treponemes as evidenced by the inter-subspecies substitutions (grey boxes) in *Treponema endemicum* (*TEN*) TP0865 proteoform (*). In contrast, TP0865 was highly conserved among SS14-lineage strains. Non-conservative substitutions are colored in red. Residues are numbered according to their position in the Nichols reference sequence. **(B)** Substitutions mapped onto the AlphaFold model for the Nichols reference. A portion of the hatch is hidden in the model to facilitate visualization

### OMP profiles reflect divergent OMP evolution within clades and subclades

We next sought to ascertain how OMP variability within our cohort correlated with the phylogenetic structure of *TPA*. We began by concatenating proteoforms for all six OMPs in each strain to generate OMP profiles (Figure 7A and Supplementary Table 3). Intriguingly, no strain within our cohort contained a profile identical to the Nichols and SS14 clade references (profiles 1^N^ and 1^S^, respectively). We identified 26 profiles among the 39 Nichols-lineage strains (profiles 2^N^ - 27^N^) compared to 24 within the 147 SS14-lineage strains (profiles 2^S^ - 25^S^). Strikingly, no OMP profiles were shared between Nichols subclades, while 27 of 30 SS14 Omega A and 14 of 24 SS14 Omega B strains shared profile 3^S^. Nichols B was the only subclade in which all strains (five from Guangzhou) contained the same profile (profile 9^N^). In a number of instances, variability of profiles within a clade can be attributed to SNVs in individual OMPs. For example, OMP profiles within the Nichols C strains contained a variable TP0548 mainly due to substitutions in the hatch, while hypervariability in ECL3 of TP0865 accounted for the high number of profiles within the Nichols E subclade. Within the SS14 clade, SNVs in ECL2 of TP0548 largely accounted for the divergent profiles within Omega C strains.

**Figure 7.**
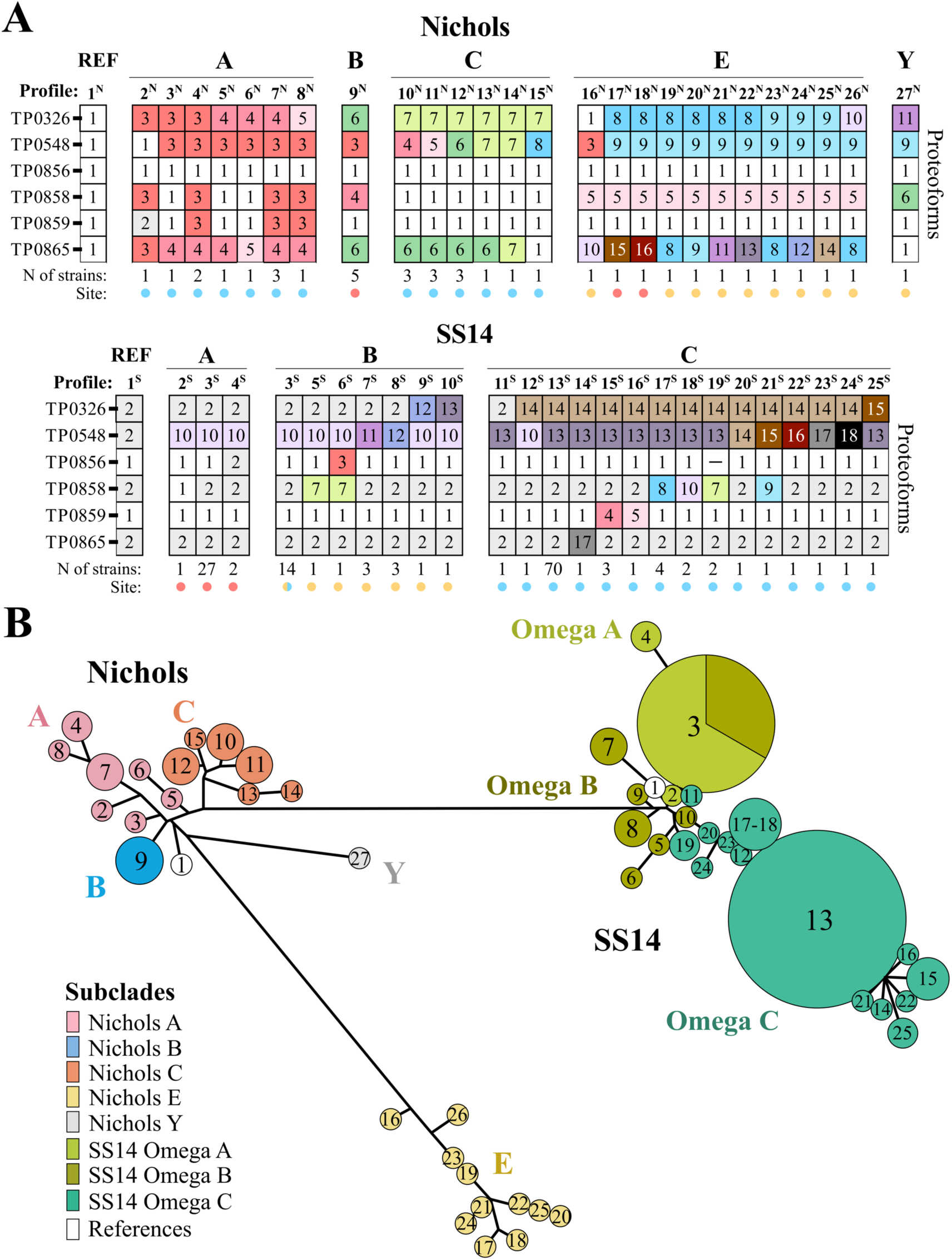
OMP profiles reflect divergent OMP evolution within clades and subclades. **(A)** OMP profiles (columns) were created by concatenating proteoforms for all six OMPs (rows) in each strain. Also shown are the number (N) of strains sharing a profile and their respective sites (colored dots: Lilongwe, blue; Guangzhou, red; Cali, yellow). **(B)** Minimum spanning tree (MST) for concatenated sequences was created using the stand-alone GrapeTree package with the neighbor-joining algorithm. Clusters are labeled with the proteoform numbers assigned in A. Strain IDs and their corresponding profiles are listed in Supplementary Table 3.

To further elucidate the relationships between OMP profiles, we next generated OMP-based minimum spanning trees (MSTs) using the GrapeTree stand-alone tool(44)(Figure 7B). The MSTs underscored that OMP profiles segregated strictly by clade, and largely by subclade, as well as the greater degree of variability within the Nichols clade. It was noteworthy that the Nichols reference profile fell on a branch separate from the other Nichols profiles, whereas the SS14 reference clustered closer with the Omega A and B profiles. The divergence of the profiles in the single Nichols Y strain and in all the Nichols E strains was mainly due to the recombinant stretches in TP0858 and TP0865, respectively. A striking difference between clades was the predominance of two profiles (3^S^ and 13^S^) within the SS14-lineage strains, whereas no predominant profile was observed within Nichols-lineage strains. The single glycine insertion in the hatch of TP0548 separated profile 3^S^ from the SS14 reference. Divergence of the Omega C profiles was mainly due to SNVs in ECL8 of BamA and ECL2 of TP0548.

### Variable and conserved surface-exposed regions of *TPA* BamA and FadLs contain predicted B cell epitopes

As a starting point for assessing the role of immune pressure in driving evolution of OMPs in *TPA*, we used the DiscoTope(45) and ElliPro servers(46) to correlate the position of predicted B cell epitopes (BCEs) with the mutations identified in the six OMPs analyzed herein. As shown in Figure 8, the vast majority of the variability occurred within regions containing predicted BCEs. A BCE is predicted for the region in BamA ECL3 (533–550) and for the hatch and ECLs in TP0548 where high inter- and intra-clade variability was found. The surface-exposed regions in TP0858, most notably ECLs 2 and 4, with intra-clade mutations, also contained predicted BCEs. Also noteworthy were the BCE predictions for TP0865 in the hypervariable ECL3. Interestingly, the algorithms also made strong epitope predictions for more conserved ECLs in BamA (ECL4) and TP0856 (ECLs 2 and 4), known to be immunogenic(17, 18, 47). Additionally, several regions (hatch, ECL3 and ECL4) of the highly conserved TP0859 also contained predicted BCEs.

**Figure 8.**
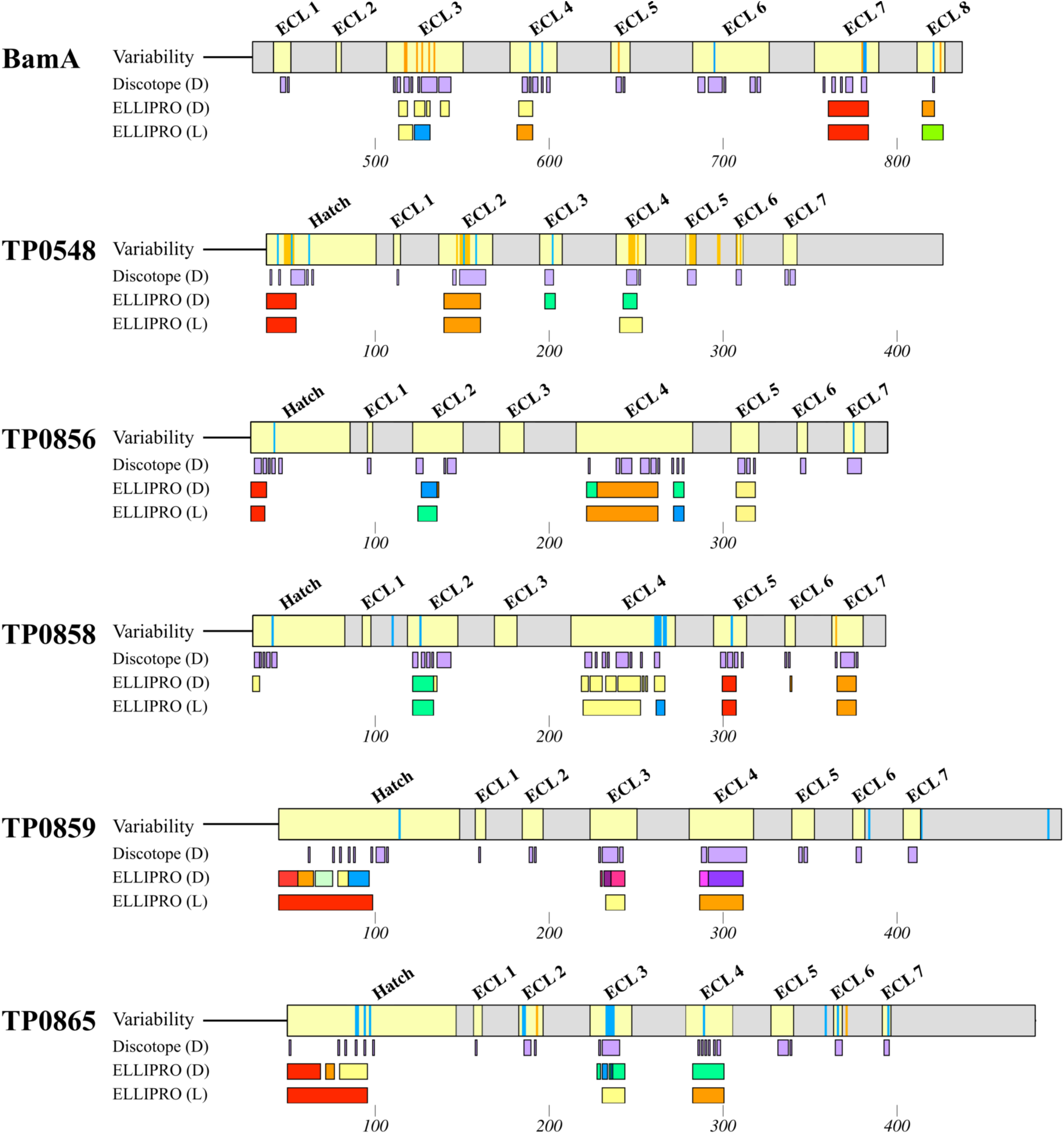
Conserved as well as variable regions of *TPA* OMPs contain predicted B-cell epitopes. DiscoTope and ELLIPRO servers predict Discontinuous (D) and Linear (L) B-cell epitopes (BCEs) for hatch and ECL regions (yellow) in all six analyzed OMPs. Position of inter-clade (orange stripes) and intra-clade (blue stripes) mutations correlate with predicted BCEs. BCEs also were predicted for conserved surface-exposed regions. Individual epitopes predicted by ELLIPRO were distinguished using different colors; DiscoTope does not differentiate individual epitopes. Graphical representations were generated using MacVector.

### Conservation of candidate ECL vaccinogens

Vaccine design in syphilis is moving towards ECL-based immunization strategies to circumvent the many difficulties inherent in the large-scale production of full-length OMP vaccinogens(7). From our studies, five ECLs (ECL4 from BamA and ECLs 2 and 4 from TP0856 and TP0858) have garnered attention as promising vaccine targets(16–19). The degree of conservation of these immunogenic ECLs, therefore, is a critical determinant of their global efficacy in a vaccine cocktail formulation(7). The Nichols and SS14 reference sequences for all five candidate ECL vaccinogens are identical, and only 16 of 186 strains in our cohort contained mutations in one or more of the five ECLs (Table 1). BamA ECL4 (1 variable strain) and TP0856 ECLs 2 and 4 (no variable strains) were highly conserved. On the other hand, candidate ECL vaccinogens from TP0858 displayed some diversity, with four SS14 Omega C strains harboring a conservative mutation in ECL2 (D141N), all 11 Nichols E strains harboring non-conservative substitutions (N277K and T281S) in ECL4, and the Nichols Y strain containing a recombination in the same ECL. Other surface-exposed regions, such as the TP0856 hatch and ECL3 and ECL4 of TP0859, contain predicted BCEs and were also highly conserved within our cohort (Figure 8), suggesting they could serve as additional targets for future vaccine development (Table 1).

**Table 1.**
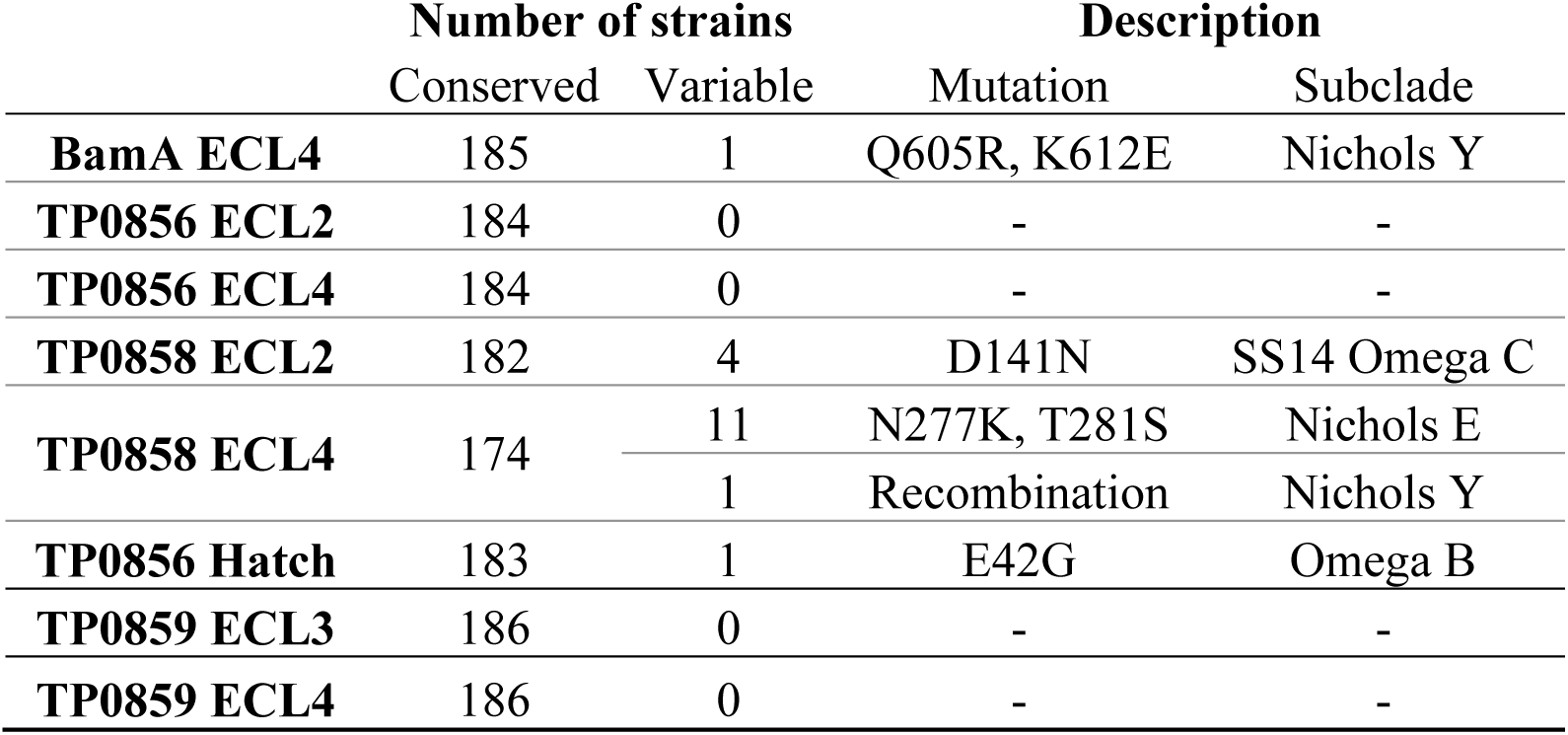
Conservation of candidate ECL vaccinogens within our cohort.

|  | Number of strains |  | Description |  |
| --- | --- | --- | --- | --- |
|  | Conserved | Variable | Mutation | Subclade |
| **BamA ECL4** | 185 | 1 | Q605R, K612E | Nichols Y |
| **TP0856 ECL2** | 184 | 0 | - | - |
| **TP0856 ECL4** | 184 | 0 | - | - |
| **TP0858 ECL2** | 182 | 4 | D141N | SS14 Omega C |
| **TP0858 ECL4** | 174 | 11 | N277K, T281S | Nichols E |
|  |  | 1 | Recombination | Nichols Y |
| **TP0856 Hatch** | 183 | 1 | E42G | Omega B |
| **TP0859 ECL3** | 186 | 0 | - | - |
| **TP0859 ECL4** | 186 | 0 | - | - |

## DISCUSSION

In addition to guiding vaccine development, the characterization of an expanded *TPA* OMPeome provides new insights into how syphilis spirochetes adapt to geographically and demographically diverse at-risk human populations. As with other pathogens(48, 49), the need to evade the host immune response likely drives *TPA* OMP adaptation, while the extent to which OMPs can adapt by sequence variation is limited by structural and/or functional constraints. The non-canonical features of *TPA* OMPs introduce additional complexity to understanding this interplay of pressures. Our syphilis vaccine consortium described a large-scale genomic analysis of *TPA* strains(31), pointing out to the existence of population-specific differences in ECLs of two *TPA* FadLs (TP0858 and TP0865). In the present study, we expanded this analysis to investigate proteoforms for six OMPs across seven geographically structured *TPA* subclades circulating in three geographically diverse sites. We confirmed that variable residues were predominantly located in surface-exposed regions, often correlating with predicted BCEs. The identification of variable (*e.g*., BamA and TP0548) and conserved (*e.g*., TP0856 and TP0859) OMPs brought to light an unexpected uneven distribution of evolutionary pressures across this portion of the OMPeome. The similarity in OMP profiles within geographically restricted subclades suggests that regional demographic factors contribute to *TPA* OMP variability.

Recombination is a major driver of OMP diversity in bacteria, occurring both within genomes (intragenomic) and between organisms (intergenomic)(50, 51). In *TPA*, the largest and most diverse OMP family – the *TPA* repeat proteins – exhibits strong evidence of both(52–54). Intragenomic rearrangements also were reported between TP0856 and TP0858(42). In a previous study of *TPA* clinical isolates(55), we found evidence of intergenomic recombination in BamA, where several Nichols- and SS14-lineage strains harbored BamA variants matching those from other *TPA* lineages, such as Mexico A. Notably, whole-genome sequencing of the Mexico A strain revealed that its BamA variant contained sequences derived from *T. pallidum* subspecies *pertenue* (*TPE*)(56). Recent studies(43, 57) indicate that most recombination events involving BamA and FadLs occurred between ancestors of the *TPA* lineage and endemic treponemes (*TPE/TEN*). In the present study, recombination among *TPA* strains was uncommon although footprints of previously described ancient intersubspecies events were discernable. For example, recombination between a Nichols E ancestor and *TPE*/*TEN*(43, 57) likely generated the highly variable TP0865 proteoforms observed in all strains within this subclade. Similarly, sequences for BamA and TP0548 were highly divergent between Nichols- and SS14-lineage strains, aligning with ancient intersubspecies recombination involving ancestors of single *TPA* clades(43, 57). In contrast, only a single recent recombination event, a modular rearrangement previously reported for this OMP(42), was identified in a TP0858 proteoform from the Nichols Y strain. The paucity(42) of recent recombination events also suggests that the TP0858 Nichols reference proteoform identified in a SS14 Omega A strain is likely the result of a SNV rather than intergenomic recombination. Intergenomic recombination requires co-infection, which might have been more common when ancient events of genetic exchange occurred among *T. pallidum* subspecies(43, 57). Evidence for co-infection has been reported in the modern era for *TPE*(58), but, surprisingly, not for *TPA*(32, 43, 57). Our analysis, as a whole, suggests that recombination involving *TPA* OMPs is rare, despite the co-circulation of Nichols and SS14 strains across all three sites, and highlights the need for broader sampling to capture these rare events.

SNVs are an important source of variability within OMPs and can significantly alter the virulence properties of bacteria, including their ability to evade host immune responses(59–65). As a stand-alone OMP(13, 21), *TPA* BamA must adapt to immune pressure while maintaining its essential function in OM biogenesis(66). The predicted BCEs for the variable regions in BamA ECL3 and ECL7 would suggest that these are the primary loops under immune pressure. However, ECL4, which was highly conserved within our cohort, also appears to be subject to immune pressure. Previously we noted that a large proportion of clinical strains from Cali, Colombia contained a substitution (L589Q) in this loop(55) that markedly diminished reactivity with ECL4 antibodies from patients infected with *TPA* strains containing the Nichols reference BamA(55). Antibodies against ECL4 of *E. coli* BamA freeze the BAM complex in the open conformation, preventing insertion of newly synthesized OMPs into the OM bilayer(67, 68); antibodies against *TPA* BamA ECL4 are bactericidal(18, 19), presumably via the same mechanism. The absence of the L589Q substitution in the present study was surprising. It is tempting to speculate that ECL4 antibodies in patients infected with *TPA* strains harboring the glutamine at position 589 in BamA conferred a selective advantage in an endemic population in which L589 predominates.

Similar to BamA, SNVs accounted for the majority of *TPA* FadL variants but with a complex pattern across the family. While TP0856 and TP0859 were highly conserved, the other FadL orthologs displayed considerable variability, with TP0548 containing the highest number of proteoforms. Additionally, the specific ECLs exhibiting the greatest variability differed for each protein: ECL4 in TP0858, ECL2 in TP0548, and ECL3 in TP0865. *TPA* FadL orthologs feature a hatch with an extended region that, as with the ECLs, differed in their degree of variability. Variability in the hatch was particularly pronounced in TP0548 but also present in TP0858 and, to a lesser extent, in TP0856 and TP0859. The variability in some of the *TPA* FadL hatches is consistent with the AlphaFold3 predictions that these regions are surface-exposed, and therefore, likely under immune pressure. In multi-gene families, functional redundancy often enables one gene to adapt while another is constrained to an essential function(69, 70). Notably, the two highly conserved FadLs, TP0856 and TP0859, belong to structurally different FadL subgroups, each containing a more sequence-divergent paralog(13). Further detailed characterization of the individual members of this family will be needed to clarify the structural factors, functional redundancies, and immunological properties driving their divergent evolutionary trajectories. While immunological pressures undoubtedly are a principal force shaping *TPA* FadL variability, we have uncovered evidence that these proteins do not fully conform to the well-established paradigm that variability within bacterial antigens correlates with antigenicity(71, 72). ECLs 2 and 4 of the most conserved FadL (TP0856) are among the most antigenic loops during rabbit and human syphilitic infection(18, 47). Why these ECLs are sequence invariant despite ostensibly intense immunologic pressure is an intriguing question. Calculations of *TPA*’s molecular clock typically are based on genomic sequences from which some OMPs, including FadLs, have been excluded(33, 57). The spectrum of variability within the *TPA* FadL family is in accord with the notion that different regions of a bacterial genome can follow distinct molecular clocks(73). As only a portion of the OMPeome was analyzed in our study, a more comprehensive investigation will be needed to fully assess how *TPA* OMP variability reflects broader molecular clock estimates.

A major finding in our study is that the OMP profiles segregated with Nichols- and SS14-lineage strains. Previous reports have called attention to the greater genomic diversity of Nichols-lineage strains compared to those in the SS14 clade(25–28, 33). The greater variability of OMP profiles within the Nichols clade represents a striking example of how OMP diversity in *TPA* mirrors genomic structure. Generation of OMP variants is generally considered an indicator of bacterial adaptability that leads to enhanced fitness in a particular environment(74–77). However, the predominance of the SS14 lineage in all three geographic sites argues that, in the case of *TPA,* greater OMP variability does not necessarily confer a selective advantage. Analysis of *TPA* OMP profiles also revealed that variability generally followed subclade-specific patterns. For example, ECL3 of BamA was hypervariable within Nichols A, while increased variability in ECL7 was observed within Nichols E. The extreme variability of TP0865 ECL3 within the Nichols E subclade, which is flanked by regions affected by ancient recombinant events, provides another example. Similarly, mutations in the TP0548 hatch domain were prevalent in several Nichols subclades, whereas ECL2 was the variable region within SS14 Omega C strains. One possible explanation for these subclade-specific OMP patterns is that they reflect regional demographic and behavioral pressures within endemic human populations. It is also possible that spontaneous mutations tolerated in one region of an OMP influence the emergence of mutations in other regions, channeling variability in subsequent generations of the same lineage. In a global phylogenic study of *TPA* strains using a database collection of 726 genomic sequences, Beale *et al.* (2021)(33) reported identical core genomes shared by strains from 14 geographically dispersed countries, suggesting that *TPA* is genetically homogenous. For reasons that are not readily apparent, *TPA* sublineages in our cohort showed a high degree of geographic restriction(31). Moreover, all but one (profile 3^S^) of the 51 OMP profiles were identified in single locations, suggesting that OMP variability in *TPA* conforms to a phylogenetic geographical structure. It is of interest to note that our minimum spanning tree analysis placed the OMP profile of the Nichols reference strain in a divergent branch from other Nichols-lineage strains, whereas the OMP profile of the SS14 reference more closely resembled those of circulating SS14-lineage strains. The contrast between clade prototypes aligns with the conclusion from the global phylogenomic analysis(33) that the Nichols reference strain represents a lineage that may no longer be widely circulating.

The surface-exposed regions of *TPA* OMPs reside at the frontline of the ‘arms race’(78) between the spirochete, an extracellular pathogen(5, 14, 15, 19), and its obligate human host. Although our study does not, strictly speaking, provide a longitudinal analysis of *TPA* OMP evolution, it offers an extended snapshot of OMP diversity that can be dissected experimentally to better understand how the spirochete ‘wages’ war within human populations. We coined the term stealth pathogenicity to describe the capacity of *TPA* to evade host antibody responses and establish persistent infection in individuals(5, 10, 11). A key question is how OMP variability promotes stealth pathogenicity within endemic populations. Experiments currently are underway to determine how amino acid substitutions affect antigenic and surface reactivity of ECL-specific antibodies; recently developed mutagenesis techniques for *TPA*(19, 79) also can be applied to solve this problem. Generating protective ECL antibodies has emerged as a major strategy for ending the arms race(16–19). Even variable *TPA* OMPs contain highly conserved ECLs, some already shown to elicit functional antibodies in rabbits and mice(16–19). Recently, we reported that immune sera from rabbits infected with either the Nichols or SS14 strain have a diminished capacity to impair the viability of the opposite clade during *in vitro* cultivation(18, 19). This observation suggests that antibodies that recognize variable ECLs may be important for complete protection and, by extension, that a broadly protective syphilis vaccine may require a ‘cocktail’ of conserved and variable ECLs.

Some limitations to our study should be acknowledged. First, by focusing on a subset of the *TPA* OMPeome, our analysis did not include OMPs that may be essential for understanding the full adaptive landscape of *TPA*. Additionally, our cohort contained strains from only three countries and almost certainly does not fully capture the global diversity of *TPA* strains. Moreover, we do not know whether some *TPA* subclades (*e.g*., Nichols Y) are truly rare in our study sites or were underrepresented due to unintended sampling biases. Nevertheless, the cardinal strength of our study is the creation of a structural framework for analyzing multiple OMPs, which, combined with deep sampling in three clinical research sites, yielded a high-resolution view of *TPA* OMP diversity within a geographic and evolutionary context. Future efforts will expand this approach to additional OMPs and sites, enabling further clarifying how host pressures at the population level shape the global *TPA* OMPeome in addition to advancing syphilis vaccine development.

## METHODS

### Study design and sample collection

We conducted a multi-center study to recruit individuals with early syphilis (primary, secondary and early latent syphilis) who presented at a sexually transmitted infection (STI) clinic in Lilongwe, Malawi; a provincial dermatology hospital in Guangzhou, China; and in a public healthcare sector network in Cali, Colombia. The recruitment period spanned from November 2019 to May 2022. Details about eligibility criteria for participant enrollment, including obtaining informed consent, are described in Sena *et al*. (2024)(31). Additional information about strains within our cohort are available in Supplementary Table 1.

### Whole genome sequencing and variant calling

Specimens containing 40 or more copies of *TPA polA* per µL underwent *TPA* enrichment followed by WGS(31). Enrichment was performed using parallel, pooled whole-genome amplification(30) and custom 120-nucleotide RNA oligonucleotide baits (Agilent Technologies, Santa Clara, CA, US). Libraries were generated using either Sure Select XT Low Input or XTHS2 kits as previously described(31). Pooled enriched libraries subsequently were sequenced using MiSeq or NovaSeq platforms (Illumina, San Diego, CA, US), employing paired-end sequencing with 150-base pair reads. Sequencing data was analyzed using a pipeline available at https://github.com/IDEELResearch/tpallidum_genomics. Briefly, adapters were removed using Trimmomatic (v0.39)(80), and host genomes were filtered out using bbmap (v38.82)(81) by mapping reads to human (hg19) or rabbit (oryCun2) reference genomes. Sequence alignments were performed with bwa (v0.7.17)(82) against the Nichols (CP004010.2) and SS14 (CP004011.1) reference sequences. Post-alignment, sequences underwent filtering to exclude reads with excessive mismatches, excessive soft or hard clipping, chimeric reads, and low mapping quality. Only sequences with at least 80% coverage by 3 reads were retained. Variant calling was conducted using GATK HaplotypeCaller (v4.4)(83), and consensus genomes were constructed with GATK FastaAlternateReferenceMaker.

### Phylogenetic analysis

For phylogenetic analysis, we used only SNPs to infer evolutionary relationships. To ensure data reliability, we masked *TPA* genomes for repetitive and difficult-to-sequence genes, including *arp*, *tp470*, and the *tpr* gene cluster. Additionally, we employed Gubbins (v3.2)(84) to identify and remove putative recombination regions. Genome alignments were performed using MAFFT (v4.790)(85) with Nichols (CP004010.2) and SS14 (CP004011.1) as reference genomes. The final phylogenetic tree was constructed with RAxML (v8.2.12)(86) and visualized using R (v4.1.2)(87) with the ggtree package (v3.2.1)(88). Clades and subclades were manually assigned to facilitate comparison with published analysis(31, 35).

### Characterization of BamA and FadL proteoforms

Sequences for BamA and FadL orthologs (TP0548, TP0856, TP0858, TP0859 and TP0865) were extracted and their proteoforms further analyzed using a Biopython script available at https://github.com/ebbettin/UCH_SRL. To avoid false variant calls, homopolymeric regions in genomic sequences from clinical strains were manually curated to match the references. Consequences of missense variants were defined using Grantham’s physicochemical distances(38), where substitutions with Grantham differences >50 were considered non-conservative. Sequences for BamA and FadLs within the same strain were concatenated and minimum spanning trees for concatenated sequences were created using the stand-alone GrapeTree package with the neighbor-joining algorithm(44).

### Structural modeling and B-cell epitope predictions

Three-dimensional models for *TPA* Nichols reference strain proteome (UP000000811) were generated using AlphaFold3(36). Protein models were visualized using UCSF Chimera v1.1(89) and identified mutations were mapped using attribute assignment files generated by the Biopython script described above. To delineate ECL boundaries, positioning of proteins in the membrane were predicted using the OPM (orientation of proteins in membranes) database(90, 91). Linear and discontinuous epitopes were predicted by ElliPro(46) and DiscoTope 3.0(45) using cut-off values of 0.7 and 1.5, respectively. Graphical representations of protein sequences were created with MacVector v18.5.

## SUPPLEMENTARY INFORMATION

**Supplementary Figure 1.** AlphaFold models for *TPA* BamA and FadLs. Protein models generated by AlphaFold3 are colored according to a per-residue confidence metric (pLDDT) ranging from 1 to 100. Regions with pLDDT values >70 (light blue) and >90 (dark blue) denote high and very high confidence backbone predictions, respectively. All β-strand residues exhibited high confidence values at a minimum, supporting accurate delineation of the boundaries between β-strands and ECLs.

**Supplementary Figure 2.** Comparison of *T. pallidum* BamA models generated by AlphaFold and ModWeb. The model for BamA generated by AlphaFold3 predicts that ECL3 (yellow) is longer and ECL4 (green) is shorter than previously predicted by the ModWeb server using the solved structure of *Neisseria meningitidis* BamA ortholog as a template. ECL numbers are shown inside circles. Figures were generated using UCSF Chimera v1.1.

Supplementary Figure 3. Inter-subspecies recombination events previously reported for *TPA* BamA and FadL. **(A)** A recombinant event in a BamA region, corresponding to the nucleotide residues 347,027 to 347,956 of the Nichols reference genome, occurred between *TEN* and the SS14 clade. **(B)** In TP0548, a recombination between *TEN* and the Nichols clade occurred between residues 593,563 and 594,215. **(C)** Two previously reported recombination events in TP0865 involving *TEN/TPE* were identified here in all Nichols E strains (nucleotide residues 945,224 to 945,542 and 945,830 to 946,298). Mutation in the hypervariable ECL3 does not reside in any of the recombinant regions.

---

**Supplementary Table 1.** Description of *TPA* strains included in the study.

**Supplementary Table 2.** Description of identified proteoforms and their respective nucleotide mutations.

**Supplementary Table 3.** Description of OMP profiles by strain.

## ACKNOWLEDGMENTS

This project was funded by the US NIH National Institute of Allergy and Infectious Diseases (NIAID; U19AI144177 to JDR and MAM) and supported in part by the Bill & Melinda Gates Foundation (INV-036560 to ACS), strategic research funds from Connecticut Children’s, NIAID (T32AI007151 to FA), and the Czech Republic National Institute of Virology and Bacteriology (Programme EXCELES LX22NPO5103; funded by the EU - Next Generation EU to DS). JAG-L was supported by a Wellcome Trust International Master’s Fellowship (214641/Z/18/Z) and a Fogarty International Center Global Infectious Diseases Research Training grant (D43TW006589). Connecticut Children’s funded the collection of samples from another longitudinal study in Colombia included in the genomic analysis of *TPA*. The authors thank all study participants and research staff in Guangzhou, Cali, Lilongwe, and Chapel Hill. We acknowledge Myron Cohen for his support, Julia Sung for participant enrollment, Carol Ospina for sample collection and management, and Santiago Camacho, Maria Fernanda Amortegui, and Nelson Romero for participant enrollment. Andreea Waltmann and Fredrick Nindo for early contributions to genomic sequencing and data analysis. Julie Nelson and Edward Jere for assisting with Malawi sample processing and analysis; Nancy Saravia provided project guidance, and Julie Vigil oversaw regulatory compliance. The authors also thank the clinical and administrative staff of private and public health institutions in Cali for referring study participants.

## AUTHOR CONTRIBUTIONS

ACS, JCS, MJC, KLH, BY, MAM and JDR conceptualized this work. ACS, JCS, MJC, KLH, and JDR organized and supervised the clinical sites. WC, FV-C, SS, JAG-L, LGR, YJ, LY, HZ, BY, MMM, IFH, and EL-M contributed to specimen acquisition and DNA extraction. FA, CMH, STH, WC, PP, DS, and KN contributed to sequencing and analysis of *TPA* genomes. EB, AAG, TCD, MJC and KLH worked on variant identification and structural mapping. EBB, AAG and JDR wrote the manuscript. All authors reviewed the manuscript.

